# Epigenomic analysis of Formalin-Fixed Paraffin-Embedded samples by CUT&Tag

**DOI:** 10.1101/2023.06.20.545743

**Authors:** Steven Henikoff, Jorja G. Henikoff, Kami Ahmad, Ronald M. Paranal, Derek H. Janssens, Zachary R. Russell, Frank Szulzewsky, Sita Kugel, Eric C. Holland

**Affiliations:** Basic Science Division, Fred Hutchinson Cancer Center, Seattle, WA, USA; Howard Hughes Medical Institute, Chevy Chase, MD, USA; Human Biology Division, Fred Hutchinson Cancer Center, Seattle, WA, USA

## Abstract

For more than a century, Formalin Fixed Paraffin Embedded (FFPE) sample preparation has been the preferred method for long-term preservation of biological material. However, the use of FFPE samples for epigenomic studies has been difficult because of chromatin damage from long exposure to high concentrations of formaldehyde. Previously, we introduced Cleavage Under Targeted Accessible Chromatin (CUTAC), an antibody-targeted chromatin accessibility mapping protocol based on CUT&Tag. Here we show that simple modifications of our single-tube CUTAC protocol are sufficient to produce high-resolution maps of paused RNA Polymerase II (RNAPII) at enhancers and promoters using FFPE samples. We find that transcriptional regulatory element differences produced by FFPE-CUTAC distinguish between mouse brain tumor specimens and identify regulatory element markers with high confidence and precision. Our simple workflow is suitable for automation, making possible affordable epigenomic profiling of archived biological samples for biomarker identification, clinical applications and retrospective studies.

## Introduction

The standard workflow of surgical specimens is from the operating room into formalin and then embedding into paraffin (FFPE), cut into sections for histological analysis and stored as paraffin blocks. Even after long-term storage, FFPE sections can be resurrected for application of modern sequencing-based genomic methodologies in ongoing and retrospective studies (1). The preferred method for archival sample preservation has been fixation in formalin (∼4% formaldehyde) for a few days followed by dehydration and embedding in paraffin. Formalin-fixed paraffin-embedded (FFPE) sample preservation has been in use for over a century, with billions of cell blocks accumulated thus far, and no end in sight (2). Most genomic studies using FFPE samples have applied whole genome sequencing to identify mutations and aneuploidies, or whole exome sequencing to identify tissue-specific differences. However, chromatin profiling has the potential of identifying causal regulatory element changes that drive disease. The prospect of applying chromatin profiling to distinguish regulatory element changes is especially attractive for translational cancer research, insofar as mis-regulation of promoters and enhancers in cancer can provide diagnostic information and may be targeted for therapy (3). However, there has been limited progress in applying chromatin profiling techniques to FFPEs (4). Although several methods have been developed for chromatin immunoprecipitation with sequencing (ChIP-seq) using FFPEs (5-10), ChIP-seq is not well-suited for small amounts of material that are typically available from patient samples. Furthermore, solubilization of such heavily cross-linked material is extremely challenging, requiring strong ionic detergents and/or proteases in addition to controlled sonication or Micrococcal Nuclease (MNase) digestion treatments.

Alternatives to ChIP-seq for chromatin profiling include ATAC-seq (11) and enzyme-tethering methods such as CUT&RUN (12) and CUT&Tag (13). Modifications to the standard ATAC-seq protocol were required to make it suitable for FFPEs, including nuclei isolation following enzymatic tissue disruption and *in vitro* transcription with T7 RNA polymerase (14, 15). The same group also similarly modified CUT&Tag and included an epitope retrieval step using ionic detergents and elevated temperatures, which they termed <u>F</u>FPE tissue with <u>A</u>ntibody-guided <u>C</u>hromatin <u>T</u>agmentation with <u>seq</u>uencing (FACT-seq) (16, 17). However, FACT-seq is a 5-day protocol even before sequencing, and the many extra steps required relative to CUT&Tag have raised concerns about experimental variability (4).

We wondered whether a fundamentally different approach to what has been described for FFPE-ATAC and FACT-seq might overcome the obstacles that have thus far been encountered in chromatin profiling of FFPEs. Rather than enzymatically breaking down the tissue for nuclei isolation, we use only heat and minimal shearing of the tissue, then follow our standard CUT&Tag-direct protocol with minor modifications. This includes applying our <u>C</u>leavage <u>U</u>nder <u>T</u>argeted <u>A</u>ccessible <u>C</u>hromatin (CUTAC) strategy, which preferentially yields <120-bp fragments released by antibody-targeted paused RNA Polymerase II (RNAPII) (18, 19). Because of the small size of the fragments released with CUTAC, it is relatively robust to the serious DNA degradation that occurs during cross-link reversal (20), and by attaching to magnetic beads and following the single-tube CUT&Tag-direct protocol we minimize experimental variation. The resulting FFPE-CUTAC profiles could be used to confidently distinguish different mouse brain tumors from one another and from normal brains, identifying potentially key regulatory elements involved in cancer progression.

To evaluate the ability of our approach to discriminate between archived samples, we chose blocks of three related mouse CNS tumor types, driven by distinct mechanisms. We compared FFPE blocks of tyrosine kinase active PDGFB-driven gliomas (21), ZFTA-RELA gene fusion-driven ependymomas (22), and YAP1-FAM118b gene fusion-driven ependymomas (23) to one another and to FFPE blocks of normal mouse brain. Analysis of FFPE-CUTAC datasets revealed that post-translational modifications marking paused RNAPII and nucleosomes at active promoters and enhancers distinguished between tumors and were elevated relative to normal at murine transcriptional regulatory elements genome-wide. We observed similar robust distinctions between mouse liver tumors and normal livers, confirming the generality of our approach.

## Results

### CUT&Tag streamlined protocol for whole cells

We originally introduced CUT&Tag with DNA purification by addition of SDS/Proteinase K followed by either phenol-chloroform-isoamyl alcohol extraction and ethanol precipitation or SPRI bead binding and elution for PCR (13). Later we streamlined the protocol so that it could be performed in single PCR tubes using a 58^°^C incubation in 0.1% SDS followed by excess Triton-X100, which sequesters the SDS in micelles, allowing efficient PCR (18). However, this CUT&Tag-direct method was only suitable for up to ∼50,000 nuclei, as more material was found to inhibit the PCR. To make CUT&Tag-direct applicable to whole cells, we have included 0.05% Triton-X100 in all buffers from antibody addition through tagmentation, which maintains cells permeable without disrupting nuclei and improves bead behavior. We have also increased the concentration of SDS and included thermolabile Proteinase K in the fragment release buffer. After digestion at 37^°^C and inactivation at 58^°^C, the SDS is quenched with excess Triton-X100 and the material is subjected to PCR, resulting in high yields with 30,000-60,000 cells (**Figure S1a-b**). When applied to the H3K4me3 promoter mark, this modified CUT&Tag-direct protocol for native whole cells resulted in representative profiles that match those of native or fixed nuclei using either the original organic extraction method or CUT&Tag-direct (**Figure 1a**). Based on MACS2 peak-calling and Fraction of Reads in Peaks (FRiP), we obtained slightly more peaks called and similar FRiP values for up to at least 100,000 native whole cells using the modified protocol (**Figure 1b-c**), obviating the need to purify nuclei for CUT&Tag-direct and AutoCUT&Tag (25).

**Figure 1:**
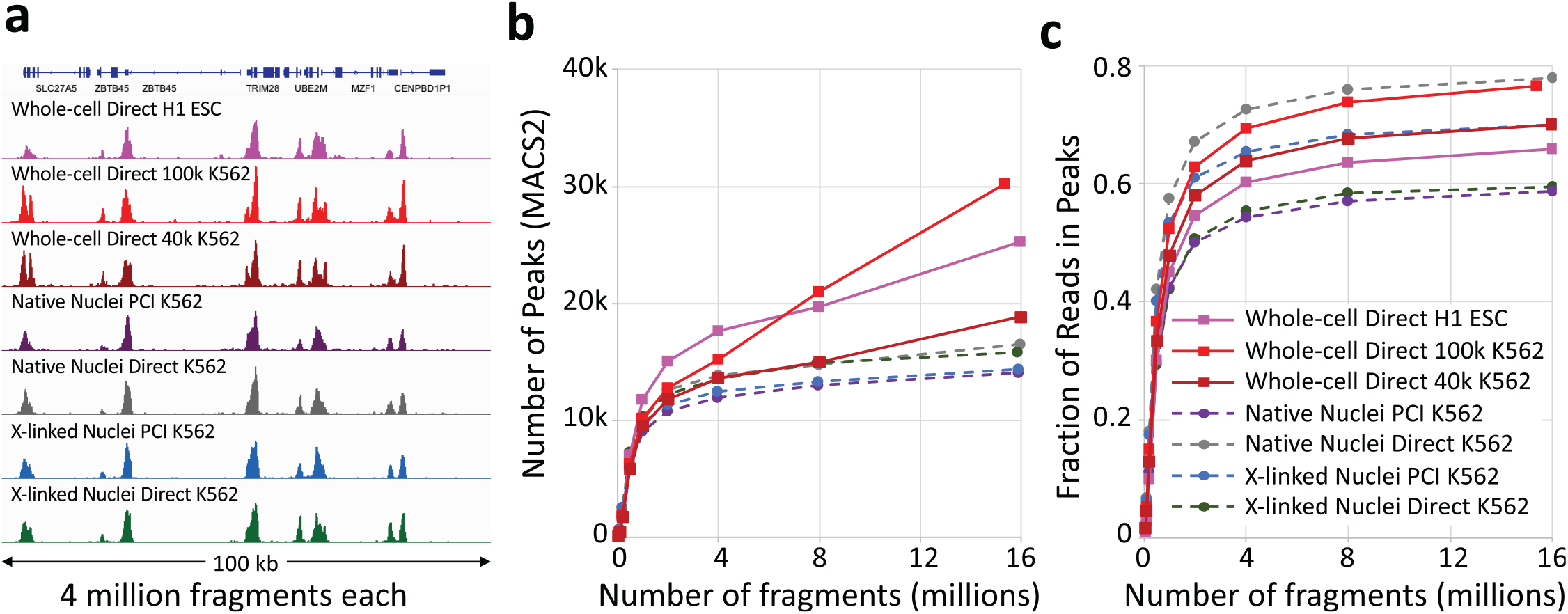
High data quality from CUT&Tag-direct for whole cells. **a)** A comparison of H3K4me3 CUT&Tag tracks for K562 cells (tracks 2-6) at a representative 100-kb region of housekeeping genes, showing group-autoscaled profiles for 4 million mapped fragments from each sample. **b-c**) Number of Peaks and Fraction of Reads in Peaks called using MACS2 on samples containing the indicated number of cells. Random samples of mapped fragments were drawn, mitochondrial reads were removed and MACS2 was used to call (narrow) peaks. The number of peaks called for each sample is a measure of sensitivity, and the fraction of reads in peaks (FRiP, right) is a measure of specificity calculated for each sampling from 50,000 to 16 million fragments. Nuclei data are from a previously described experiment (55).

### Temperature-dependent permeabilization of FFPE sections for CUTAC

The difficulty of performing CUT&Tag-direct on FFPEs is exacerbated not only by the severe chromatin damage caused by heavy formalin fixation but also by the large amount of cross-linked intra- and inter-cellular material that cells are embedded in. Both the FFPE-ATAC and FACT-seq methods require lengthy digestion with collagenases and hyaluronidase followed by 27-gauge needle extraction and straining liberated nuclei for processing. We reasoned that harsh treatments might not be necessary if the cells can be permeabilized sufficiently, and we were encouraged to attempt this approach by the fact that deparaffinized 5-10 micron FFPE samples on slides are routinely permeabilized for cytological staining with antibodies (1). Also, there has been recent progress in preventing the most severe DNA damage to FFPEs by careful attention to buffer and heating conditions (20). Accordingly, we performed manual shearing of deparaffinized 10-micron FFPE sections from tumor and normal mouse brains by dicing and scraping the tissue off slides with a razor blade followed by forcing the solution twenty times through a 22-gauge needle. We found that the Concanavalin A (ConA) beads used for standard CUT&Tag bound sufficiently well to sheared FFPE fragments regardless of whether they had been prepared from samples deparaffinized using a xylene or a mineral oil procedure. This meant that all steps from antibody addition through to PCR could be performed on FFPEs following the same CUT&Tag-direct protocol used for nuclei and whole cells. In addition, the toughness of FFPE shards allowed for hard vortexing and centrifugation steps that would have resulted in lysis of ConA bead-bound cells or nuclei.

Formaldehyde cross-links are reversed by incubation at elevated temperatures. Typical ChIP-seq, CUT&RUN and CUT&Tag protocols recommend cross-link reversal at 65^°^C overnight in the presence of Proteinase K and SDS to simultaneously reverse cross-links and deproteinize. However, the much more extreme formaldehyde treatments that are used in preparing FFPEs have required incubation temperatures as high as 90^°^C for isolation of PCR-amplifiable DNA for whole-genome sequencing (20, 27, 28). High temperatures also contribute to epitope retrieval for ChIP-seq (5-10) and FACT-seq (16), and for cytological staining one protocol calls for epitope retrieval at 125^°^C at 25 psi in a pressure cooker (29). To optimize the temperature of incubation for DNA recovery and epitope retrieval for CUTAC on FFPE samples from mouse brain tumors, we incubated sheared FFPEs at temperatures ranging from 65^°^C to 95^°^C before ConA bead and antibody additions. We performed modified CUT&Tag-direct using low-salt tagmentation (CUTAC) with RNAPII-Ser5p and/or RNAPII-Ser2,5p and H3K27ac antibodies. Upon DNA sequencing, the fraction of fragments that mapped to the mouse genome showed a strong temperature dependence, where the highest temperatures (90-95^°^C) showed the highest fraction mapping to the mouse genome (75%), and the lowest temperatures (65-70^°^C) showed the lowest fraction (13%) (**Figure 2a**). A relationship between cross-link reversal and incubation temperature has been determined to follow the Arrhenius equation (26). As temperature dependence of mouse tagmented fragment recovery also followed the Arrhenius equation, cross-link reversal is likely limiting for fragment recovery.

**Figure 2:**
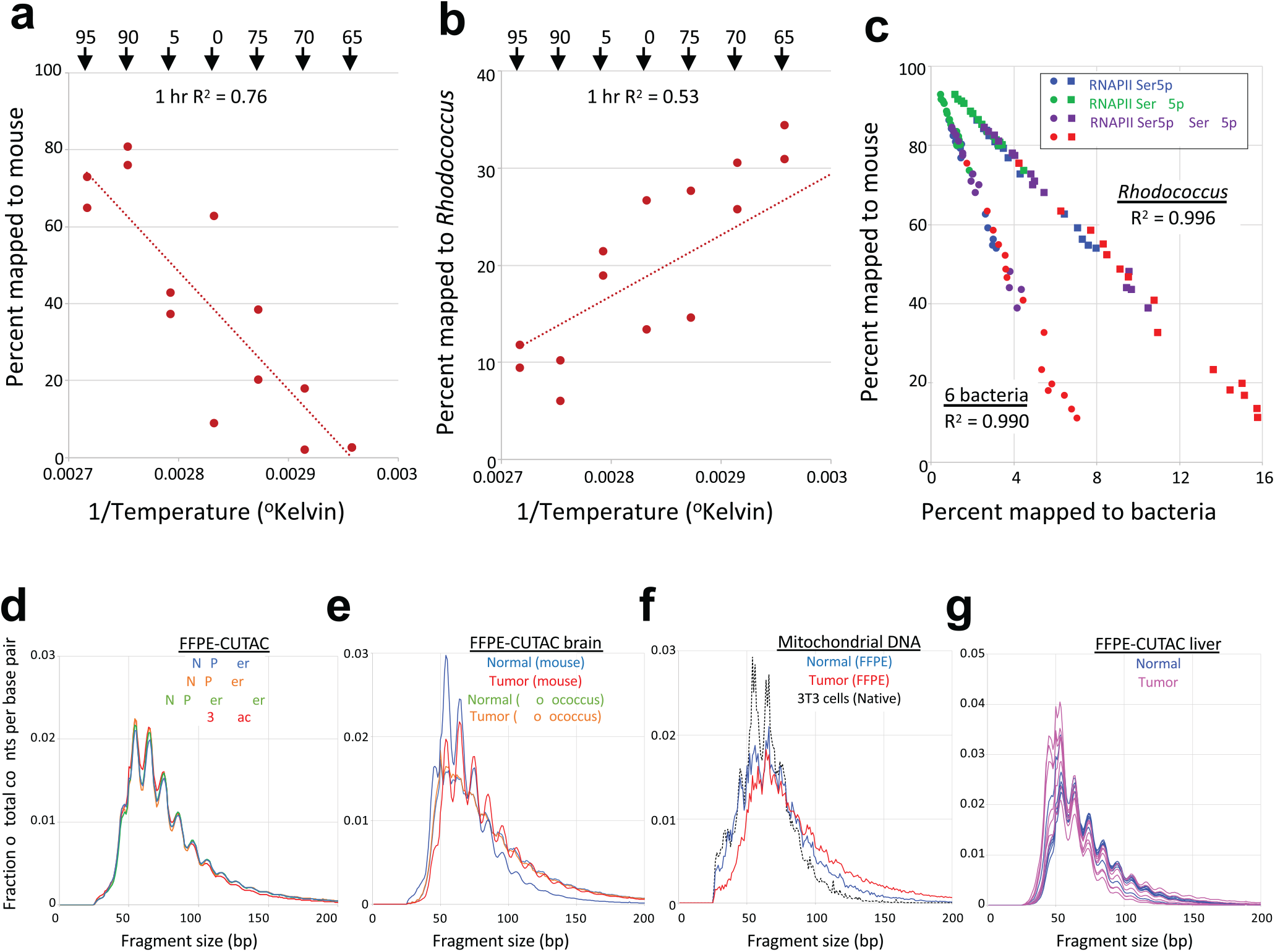
High temperatures improve yield of small mouse fragments with FFPE-CUTAC. **a)** Arrhenius plot showing the recovery of fragments mapping to the Mm10 build of the mouse genome as a function of temperature. Deparaffinized FFPEs were scraped into cross-link reversal buffer (20) containing 0.05% Triton-X100, needle-extracted, and divided into PCR tubes for incubation in a thermocycler at the indicated temperatures. **b**) Same as (a) except for fragments mapping to the *Rhodococcus erythropolis* genome. **c**) Scatter plots and R^2^ correlations between total fragments recovered versus *R. erythropolis* and the summed totals for 6 other bacterial species discovered in BLASTN searches of unmapped reads (*Escherichia coli, Leifsonia* species, *Deinococcus aestuarii, Mycobacterium syngnathidarum, Vibrio vulnificus*, and *Bacillus pumilus*). **d**) Average length distributions for three single antibodies (RNAPII-Ser5p: 15 samples; RNAPII-Ser2,5p: 15 samples; H3K27ac: 15 samples) and a 50:50 mixture of RNAPII-Ser5 and RNAPII-Ser2,5p: 14 samples. For each sample, mouse and *Rhodococcus* fragment lengths were divided by the total number of fragments before averaging. Lengths are plotted at single base-pair resolution. **e**) Average length distributions for the same samples as in (d) except grouped by cancer driver transgene (YAP1: 23 samples; PDGFB: 8 samples; RELA: 8 samples) and Normal brain: 20 samples. **g**) Same as (d) except for Mm10 ChrM (mitochondrial) fragments from the same FFPEs as used for panels e and f. The length distribution of Mm10 ChrM fragments from mouse 3T3 cells is plotted for reference. **f**) Same as (d) except that individual curves for liver tumors (magenta, 7 samples) and normal livers (blue, 6 samples) are superimposed.

### High temperatures preferentially reduce tagmentation of contaminating bacterial DNA

We were curious as to the identity of fragments generated by FFPE-CUTAC that did not map to the mouse genome. Using BLASTN against nucleotide sequences in Genbank it became apparent that there was a single species that consistently rose to the top of the list for all samples, the gram-positive bacterium *Rhodococcus erythropolis*. Mapping fragments to the *R. erythropolis* genome, we found that the entire genome was represented as expected if this species is a major contaminant of the mouse brain FFPEs in our study. Consistent with this interpretation, we found a high-temperature dependence of fragment release opposite that for mouse (**Figure 2b**), consistent with *Rhodococcu*s fragments competing with mouse fragments in PCR. We also found a near-perfect anti-correlation between the fraction of fragments mapped to mouse and the fraction mapped to the *R. erythropolis* genome (R^2^ = 0.996, n=59) across all antibodies (**Figure 2c**), with *Rhodococcus* accounting for 1-15% of the total fragments. As bacterial DNA is not chromatinized, it is unlikely to be protected from melting as well as mouse DNA, and so would not serve as a substrate for Tn5 tagmentation, which could account for the reduction in *Rhodococcus* contamination with increasing temperature.

To obtain a broader representation of species contaminating our FFPEs, we performed BLASTN searches of the RefGene Genome Database using a sample of 300 multiply represented 50-bp reads not aligning to the Mm10 build of the mouse genome. A search of the bacterial genome subset returned hits to diverse species for 208 species for ∼2/3^rd^ of the fragments, which implies that most of the unmapped reads were bacterial in origin. Although no other bacterial species were nearly as abundant as *R. erythropolis*, summing the fragment counts mapped to the six most frequently represented other species accounted for ∼0.5-7% of the fragments and showed similar near-perfect anti-correlations to mouse (R^2^ = 0.990, **Figure 2c**). Efficiency was highest for RNAPII Ser2,5p (85% mouse, 2.5% *Rhodococcus*) and lowest for H3K27ac (38% mouse, 11% *Rhodococcus*). The lower efficiency of the histone modification, and our observation that this protocol failed for H3K4me2 and H3K4me3, suggests that lysine-rich histone tails are more subject to formalde-hyde adduct and cross-linking damage than the C-terminal domain of Rpb1, which consists of 52 copies of a lysine-free YSPTSPS heptamer.

What is the source of *Rhodococcus* and other bacterial contamination in our FFPEs, which derive from multiple FFPE sample preparations over a 2-year span? *R. erythropolis* isolates have been found to use paraffin wax as their sole carbon source, forming thick biofilms (30). The species has also been proposed as an industrial biodegrader for removing the paraffin wax that remains on the inner surfaces of oil tanker holds after they are emptied (31). We infer that most of the DNA fragments that do not map to mouse are derived from the paraffin used in embedding, with an advantage during PCR over the tissue derived DNA in not having been subjected to formalin treatment. We interpret the near-perfect anti-correlations seen for these genomes in different samples as reflecting a very uniform distribution of contamination for slides prepared at different times.

### Subnucleosomal fragment sizes from FFPE-CUTAC samples

Capillary gel profiles of FFPE-CUTAC libraries revealed insert sizes averaging ∼60 bp (**Figure S1c**), despite inclusion of a 1 minute 72^°^C PCR extension step in each PCR cycle intended to capture larger fragments from degraded template DNA. After DNA sequencing, we observed subnucleosomal length distributions showing 10-bp periodicities typical of CUT&Tag peaking at ∼60 bp for all antibody series (**Figure 2d**). When we separately plotted the fragment length distributions for tumors and normal brains, we observed a conspicuous difference, where the length distribution was shifted with more longer fragments in tumors (median = 76 bp) relative to normal brains (median = 65 bp) (**Figure 2e**). In contrast, the two overall length distributions of *Rhodococcus* DNA fragments from the same tumor and normal samples closely superimposed. This shift to a longer fragment distribution for tumors is also seen for mitochondrial DNA from the same samples when compared to either normal brain or CUT&Tag mitochondrial DNA profiles from native 3T3 fibroblasts (**Figure 2f**). However, a small difference in the opposite direction was observed between liver tumor (median = 63 bp) and normal (median = 68 bp) FFPEs (**Figure 2g**), which suggests that the length differences seen between tumor and normal mouse brain are tissue-specific. Interestingly, both *Rhodococcus* and mouse mitochondrial fragments from FFPEs displayed a much weaker 10-bp periodicity relative to mouse brain FFPE nuclear and unfixed mouse mitochondrial fragments, respectively (**Figure 2f**), suggesting that the reduction in periodicity seen for DNA unimpeded by nucleosomes (bacterial and mitochondrial) is the result of DNA damage caused by fixation and cross-link reversal. The strong periodicity seen for mouse CUT&Tag profiles relative to non-chromatinized DNA of bacteria and mitochondria in the same samples might reflect partial protection from unreversible formadehyde fixation damage by RNAPII and other chromatin regulatory complexes characteristic of open chromatin (32).

### FFPE-CUTAC produces high-quality maps of active chromatin

To evaluate the accuracy and data quality of FFPE-CUTAC applied to mouse brain tumors, we compared tracks between FFPE-CUTAC and FACT-seq or standard CUT&Tag from the same study (16) using the same H3K27ac antibody (Abcam cat. no. 4729). Because of differences in cell types, brain tumors in our study and kidney or liver in the FACT-seq study, we limited comparisons of tracks to housekeeping genes that are expected to be similarly expressed in all cell types. Based on visual inspection of tracks from representative regions of the mouse genome, it is evident that H3K27ac CUTAC profiles show much cleaner profiles than those obtained using FACT-seq, with higher sensitivity than the data obtained for CUT&Tag controls of frozen mouse kidney (**Figure 3a-d**). Likewise, clean profiles were also seen for RNAPII-Ser2,5p FFPE-CUTAC, where RNAPII-Ser2 phosphate marks elongating and RNAPII-Ser5 phosphate marks paused RNAPII.

**Figure 3:**
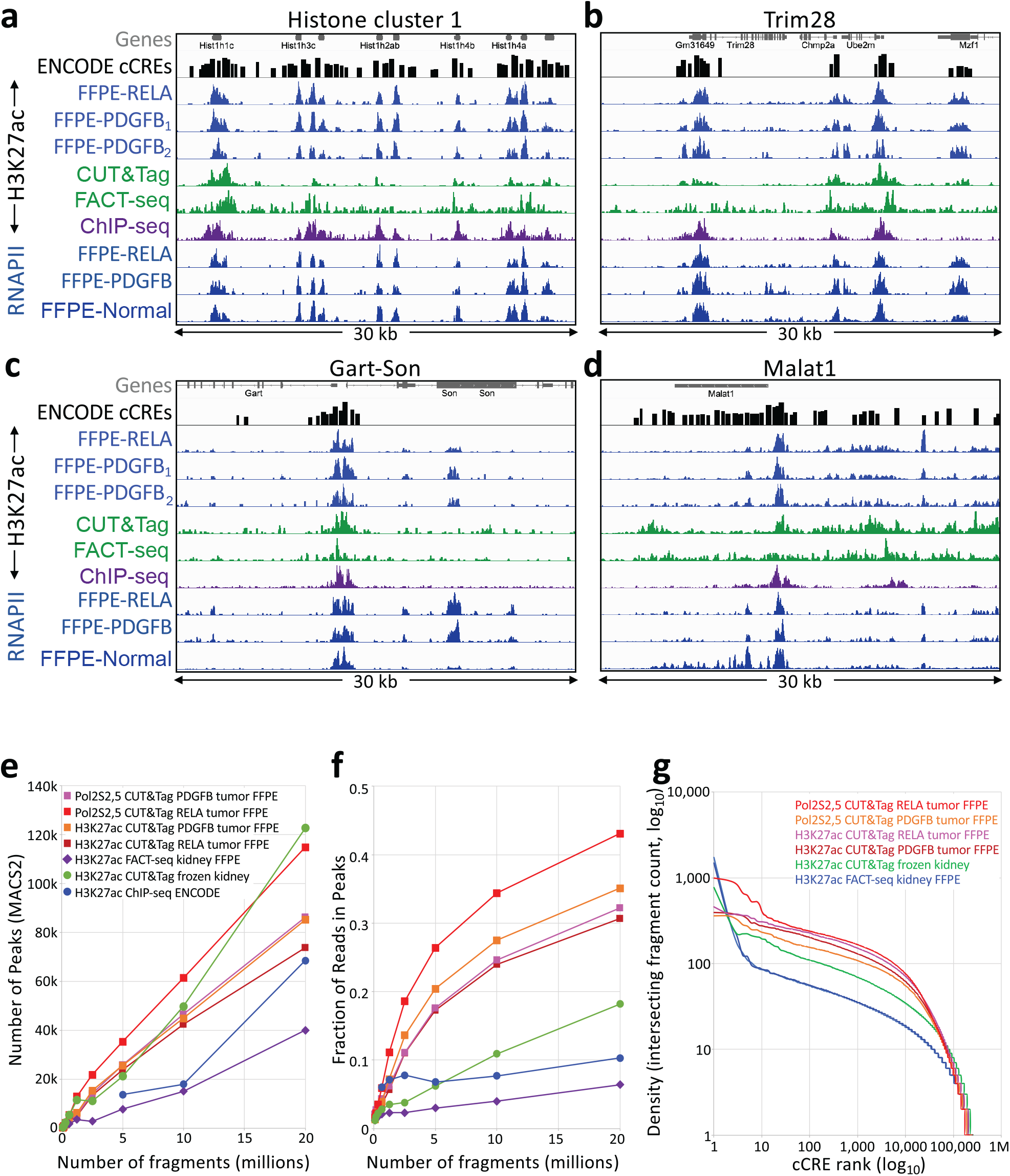
Comparison of H3K27ac FFPE-CUTAC to FACT-seq and CUT&Tag of frozen unfixed samples. Representative examples of housekeeping gene regions were chosen to minimize the effect of cell-type differences between FFPE-CUTAC (three brain tumors) and published FACT-seq and control CUT&Tag data (kidney). Forebrain H3K27ac ChIP-seq and ATAC-seq samples from the ENCODE project are shown for comparison, using the same number of fragments (20 million) for each sample. Also shown are tracks from FFPE-CUTAC samples using an antibody to RNAPII-Ser2,5p. A track for Candidate *cis*-Regulatory Elements (cCREs) from the ENCODE project is shown above the data tracks, which are autoscaled for clarity. (**e-f**) Number of peaks and Fraction of Reads in Peaks (FRiP) called using MACS2 on samples containing the indicated number of cells. **g**) Cumulative log_10_ plots of normalized counts intersecting cCREs versus log_10_ rank.

For a systematic analysis of data quality, we called peaks using MACS2 and compared the number of peaks called and FRiP values. Both H3K27ac and RNAPII-Ser2,5p FFPE-CUTAC on RELA- and PDG-FB-driven brain tumors showed much better sensitivity based on number of peaks called and much higher FRiP values than either H3K27ac CUT&Tag on frozen kidney or FACT-seq on FFPEs (**Figure 3e-f**).

To determine the degree to which FFPE-CUTAC profiles capture regulatory elements, we took advantage of the Candidate *cis*-Regulatory Elements (cCRE) database generated by the ENCODE project, which called putative regulatory elements from all tissue types profiled. We used the 343,731 elements in the cCRE mouse database based mostly on DNAseI-seq, but also H3K4me3 and CTCF ChIP-seq. This resource provides a comprehensive standard for FFPE-CUTAC performance, insofar as CUTAC profiles correspond closely to both ATAC-seq and DNAseI-seq profiles (18). For each dataset we rank-ordered cCREs based on normalized counts spanned by each element, which we plotted as a log-log cumulative curve, where a higher curve indicates better performance in distinguishing annotated sites from background. By this benchmark, both H3K27ac and RNAPII-Ser2,5p FFPE-CUTAC brain datasets outperformed both FACT-seq on FFPEs and CUT&Tag on unfixed frozen kidney (**Figure 3g**). We conclude that our FFPE-CUTAC protocol provides high quality data, even when compared to standard CUT&Tag.

### FFPE-CUTAC profiles distinguish brain tumors and reveal global upregulation

Nearly all strong peaks seen for H3K27ac and RNAPII-Ser2,5p FFPE-CUTAC corresponded to putative regulatory elements from the cCRE database, with concordance between FFPE-CUTAC, FACT-seq and ChIP-seq (**Figure 3a-d**). To identify tumor-specific candidate regulatory elements we performed pairwise comparisons between three different mouse brain tumors (YAP1-, PDGFB- and RELA-driven tumors) and normal mouse brains. For each of the 343,731 cCREs we averaged the normalized counts spanned by the cCRE and performed pairwise comparisons over all cCREs with Voom (33), an Empirical Bayes algorithm, which uses the other datasets as pseudo-replicates to increase statistical confidence. We applied this approach to datasets from multiple FFPE-CUTAC experiments using antibodies against RNAPII-Ser5p, RNAPII-Ser2,5p and H3K27ac. We observed far more significant differences for comparisons between tumors and normal brains than between tumors, with more increases than decreases in tumors relative to normal brains (**Figure 4a-c** and **Table S1a-d**). For example, using RNAPII-Ser5p, there were 10,321 cCREs that differed between YAP1 and normal brain, 518 between PDGFB and normal brain, and 190 between RELA and normal brain at a False Discovery Rate (FDR) = 0.05, but only 10-63 cCREs that differed in pairwise comparisons between the three tumors (**Figure 4a** and **Table S1a**). Compared to normal brain, 92-99% of the differences were increases in the tumors. Approximately similar results were obtained using RNAPII-Ser5p (**Figure 4b** and **Table S1b**). For H3K27ac, the number of cCREs that increased was more extreme, with nearly half of the 343,371 cCREs significantly increased at the FDR = 0.05 level **Figure 4c** and **Table S1d**). These results demonstrate that FFPE-CUTAC using RNAPII or H3K27 marks distinguishes between the tumors and the normal brain samples with nearly all significant differences representing increases for the three tumors over normal brain.

**Figure 4:**
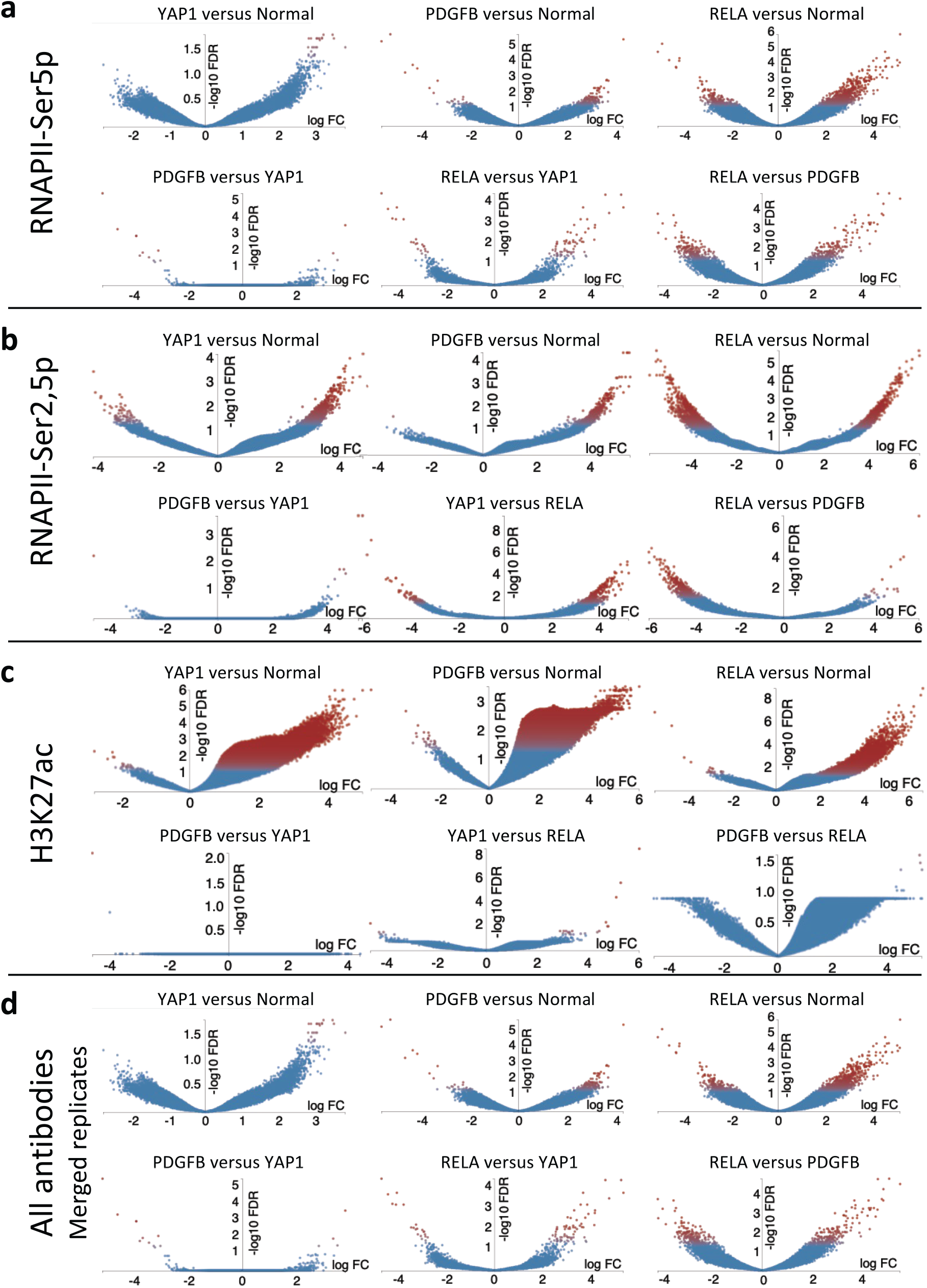
Volcano plots for pairwise comparisons between FFPE-CUTAC samples. The Degust server (https://degust.erc.monash.edu/) was used with Voom/Limma defaults to generate volcano plots, where replicates consisted of a mix of samples run in parallel or on different days on FFPE slides from 8 different brain samples (3 Normal, 3 YAP1, 1 PDGFB, 1 RELA). **a**) Comparisons based on RNAPII-Ser5p using average normalized counts per base-pair for each cCRE, applying the Empirical Bayes Voom/Limma algorithm for pairwise comparisons using the other datasets as pseudo-replicates to increase statistical power. Replicate numbers: Normal: 13; YAP1: 14, PDGFB: 3; RELA: 2. **b**) Same as (a) for RNAPII-Ser2,5p. Replicate numbers: Normal: 5; YAP1: 6; PDGFB: 3; RELA: 3. **c**) Same as (a) for H3K27ac. Replicate numbers: Normal: 10; YAP1: 12; PDGFB: 5; RELA: 7. **d**) Datasets from multiple FFPE-CUTAC experiments for each antibody (RNAPII-Ser5p, RNAPII-Ser2,5p or H3K27ac) or antibody combination (RNAPII-Ser5p + RNAPII-Ser2,5p) were merged, then down-sampled to the same number of mapped fragments for each genotype. These 16 datasets (4 antibodies x 4 genotypes) were compared against each other with Voom using the other 14 datasets as pseudo-replicates.

As FFPE-CUTAC data quality is very similar between RNAPII-Ser2,5p and H3K27ac (**Figure 3**), we attribute the conspicuous sensitivity differences in pairwise comparisons (**Figure 4a-c** and **Table S1a-d)** in part to the larger number of H3K27ac samples that Voom used for pseudo-replicates in calculating FDR. To balance the contribution of samples from each genotype, we merged datasets from multiple FFPE-CUTAC experiments for each antibody (RNAPII-Ser5p, RNAPII-Ser2,5p or H3K27ac) or antibody combination (RNAPII-Ser5p + RNAPII-Ser2,5p), then down-sampled to the same number of mapped fragments for each genotype. The three tumor and one normal genotypes, each represented by four different antibodies or antibody combination, were compared pairwise with Voom. We observed the most differences between RELA and Normal (1657) and between RELA and PDGFB (607) and the fewest differences between PDGFB and YAP1 (15) (**Figure 4d** and **Table S1e**). We conclude that FFPE-CUTAC can distinguish tumors from one another and from normal brains based on differences in cCRE occupancy of active RNAPII and H3K27ac marks.

### Increases in paused RNAPII pinpoint regulatory element differences

To identify gene regulatory elements genome-wide that best distinguish tumor from normal and between tumors, we performed Voom analysis using the maximum within each cCRE, rather than the average over the entire cCRE. The most significant difference among all RNAPII-Ser5p cCRE comparisons is a sharp peak in a coding exon of the PDGFB gene, which is present in the PDGFB-driven tumors but absent in the normal brain (FDR = 5 × 10^−5^, **Figure 5a**). This example serves as an internal control, as it corresponds to the virally expressed PDGF-beta growth factor coding region that drives the tumor, even though this sample contained both normal brain and tumorous tissue. The other most significant and highly expressed differences between tumors and normal brain identify loci that have been reported as implicated in tumor progression. Among these are the SET domain-containing 5 (*Setd5*) promoter (**Figure 5b**)(34), the Phosphoglucokinase (*Pgk1*) promoter (**Figure 5c**) (35), which are also from the PDGFB-driven tumor and normal comparison, displaying clear differences between the tumors. Additionally, the cCREs in these genes show high signal in the RELA-driven tumor and low signal in the YAP1-fusion-driven tumor. Even more striking differences are seen for the next two most significant differences at the bidirectional promoter of the Insulin growth factor 2 (*Igf2*) (**Figure 5d**) and the Collagen type 1 alpha 1 (*col1a1*) gene promoter (**Figure 5e**) (36 (37), where the RELA-driven tumor shows a strong signal but there is no perceptible signal in the region for normal, PDG-FB-driven and YAP1-driven samples. Conspicuous tumor-specific differences are also seen for four of the five cCREs with the highest signals with FDR < 0.05, including an intronic enhancer in the Suppressor of cytosine signaling 3 (*Socs3*) gene (**Figure 5f**) (38), the promoter of the Nuclear paraspeckle assembly transcript 1 (*Neat1*) long non-coding RNA gene (**Figure 5g**) (39), a proximal enhancer of the Cyclin D1 (*Ccnd1*) gene (**Figure 5h**) (40) and the C/EBPβ promoter (**Figure 5j**) (41). Additional genes implicated in tumor progression are highlighted by these comparisons, including the Connective tissue growth factor (*Ctgf*) promoter (**Figure 5k**) (42) and an intronic enhancer of the Metallothionien 2A (*Mt2a*) gene (**Figure 5l**) (42). Finally, whereas the Testis Expressed 14 (*Tex14*) gene has not been reported to be implicated in cancer, this is the only one of the top 12 genes in which the tumor/normal differences were inconspicuous (**Figure 5i**), consistent with the supposition that increases in paused RNAPII at enhancers or promoters of the other 11 genes are associated with tumor progression.

**Figure 5.**
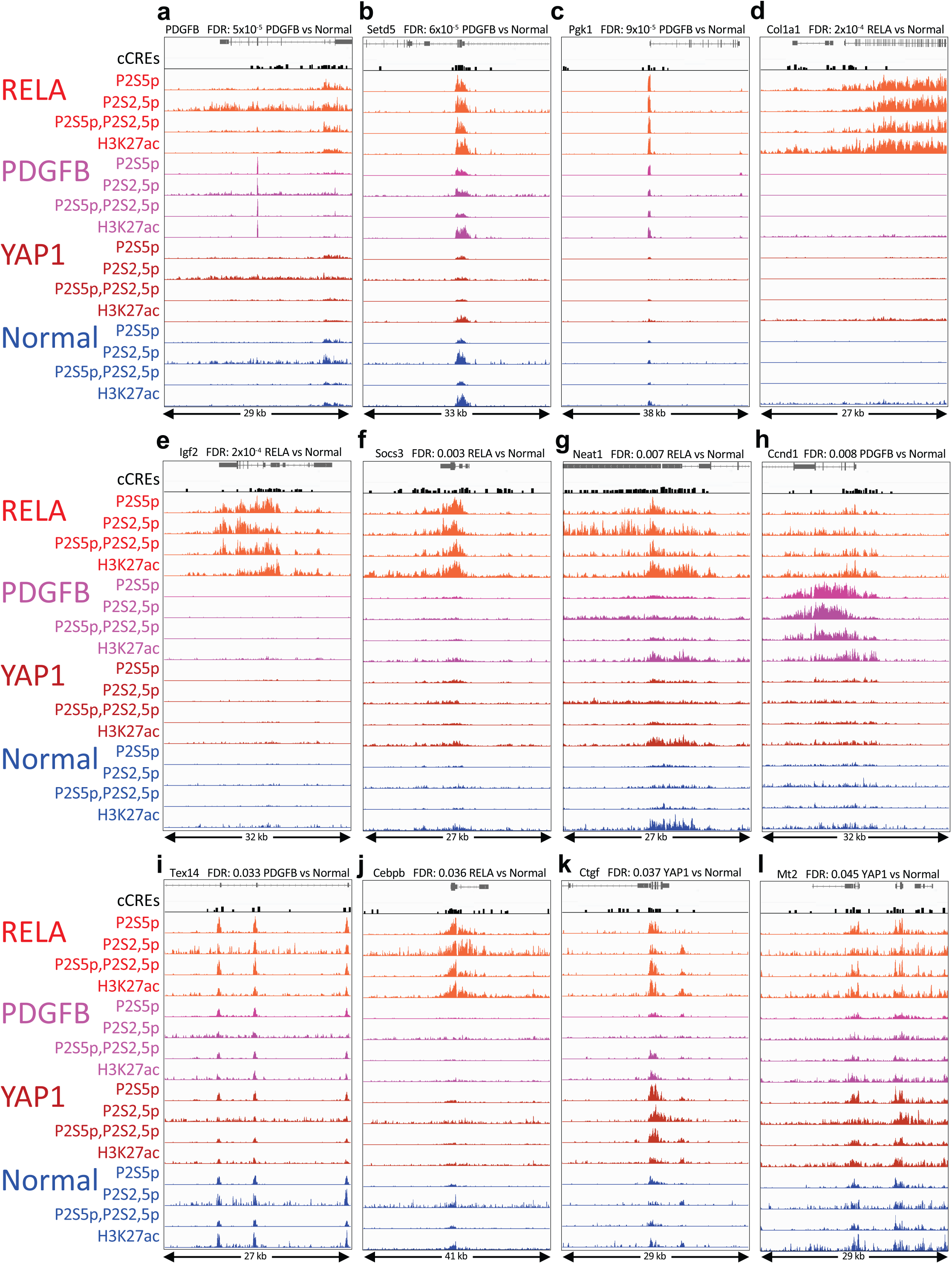
*(from previous page)*: a-e) Top significant differences between tumor and normal and between tumors based on RNAPII-Ser5p FFPE-CUTAC comparisons. IGV tracks centered around the cCREs with the most significant difference across all pairwise comparisons (FDR = 5 × 10^−5^ – 2 × 10^−4^). To enrich for regulatory elements within the span of each cCRE, we used the maximum value. **f-j**) Tracks centered around the cCRE for each of the strongest signals with FDRs < 0.05, ordered by increasing FDR (0.003 – 0.045).

### FFPE-CUTAC distinguishes tumors from normal liver

To test whether our results with mouse brain FFPEs generalize to a very different tissue type, we performed FFPE-CUTAC using FFPE sections prepared from intrahepatic cholangiocarcinoma tumors and normal liver. We used FFPE sections that had been fixed in formalin for 7 days and after deparaffinization were incubated at 90^°^C in cross-link reversal buffer for 8 hours and incubated with a 50:50 mixture of RNAPII-Ser5p and RNAPII-Ser2,5p antibodies, each at 1:50 concentration. Highly consistent results were obtained for samples ranging from 10% to 50% of a section (∼30,000-150,000 cells), with clean peaks over house-keeping genes for both liver tumor and normal liver (**Figure 6a-d**). As was the case with brain tumor and normal tissues fixed in formalin for 2 days, the number of peaks and fraction of reads in peaks (FRiP) were much higher than those from FACT-seq FFPE livers (**Figure 6e-f**), and overlap with cCREs was also much higher when down-sampled to the same number of fragments (**Figure 6g**). Finally, volcano plots revealed net increases in cCRE RNAPII occupancy both in fold-change and FDR for liver tumors relative to normal livers, similar to what we observed in comparing brain tumors to normal brains (**Figure 6h-i**). We conclude that FFPE-CUTAC provides high-quality for FFPEs from diverse tissue types.

**Figure 6:**
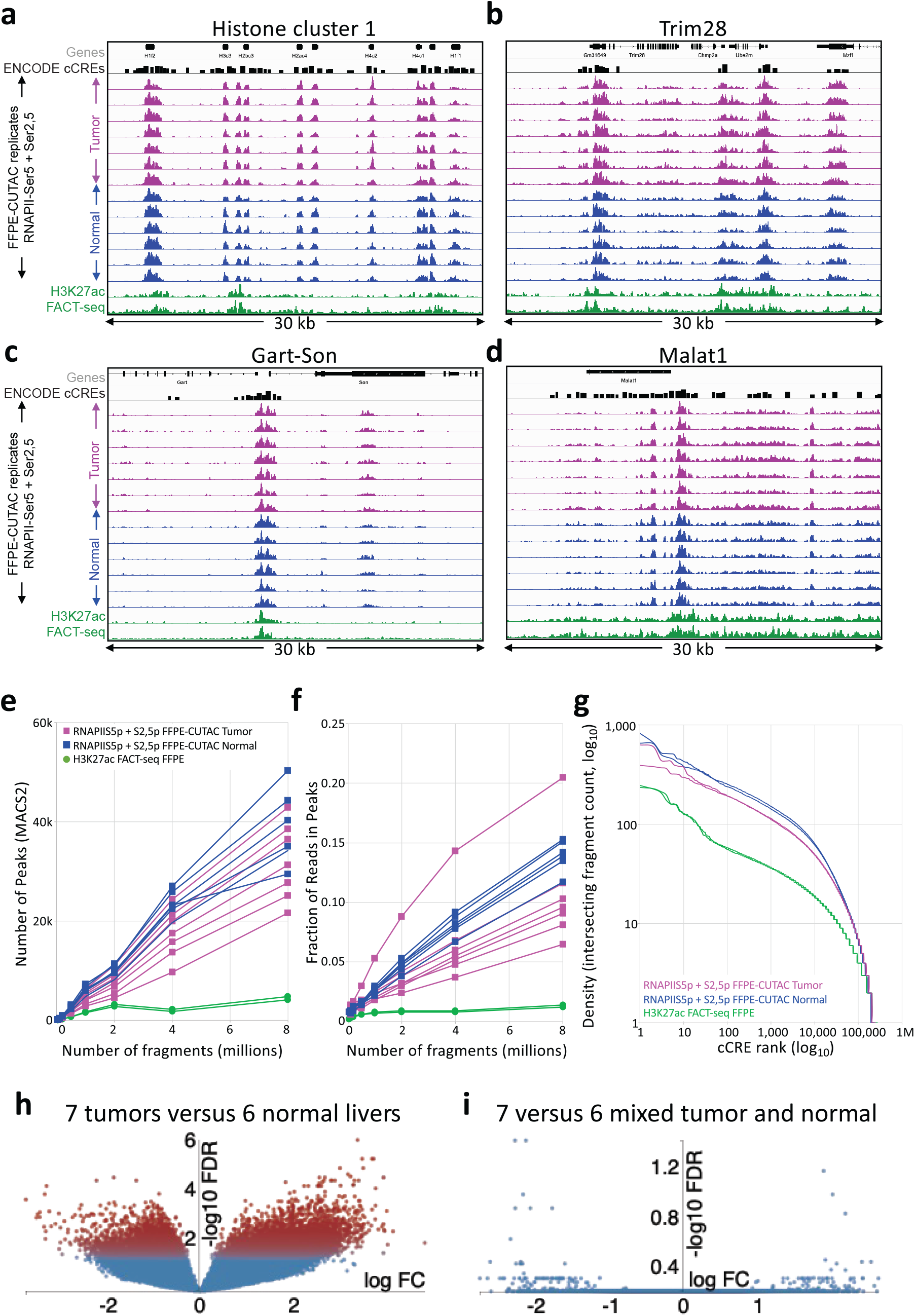
FFPE-CUTAC produces high-quality data from liver FFPEs. **a-d**) Representative tracks of liver tumor and normal liver FFPE-CUTAC and FACT-seq samples at the housekeeping gene regions depicted in Figure 3. A track for Candidate *cis*-Regulatory Elements (cCREs) from the ENCODE project is shown above the data tracks, which are autoscaled for clarity. (**e-f**) Number of peaks and Fraction of Reads in Peaks (FRiP) called using MACS2 on samples containing the indicated number of cells for 7 liver tumor (magenta), 6 normal liver (blue) and 2 normal liver FACT-seq (green) samples. **g**) Cumulative log_10_ plots of normalized counts intersecting cCREs versus log_10_ rank for representative liver samples. **h**) Voom/Limma volcano plot for the 7 liver tumor versus 6 normal liver samples. **i**) Control volcano plot in which three liver tumor samples and 3 normal livers were exchanged for Voom/Limma analysis.

### Comparison between FFPE-CUTAC and standard RNA-seq on transgene-driven brain tumors

The murine brain tumors that we used in our study have served as models for the study of *de novo* tumorigenesis (22, 23, 43), with high-quality RNA-seq data available. To do an unbiased comparison between FFPE-CUTAC regulatory elements and processed transcripts mapped by RNA-seq, we first determined whether there is sufficient overlap between cCREs and annotated 5’-to-3’ genes to fairly compare these very different modalities. Specifically, the 343,731 cCREs average 272 bp in length, accounting for 3.4% of the Mm10 build of the mouse genome, whereas the 23,551 genes in RefGene average 49,602 bp in length, with an overlap of 54,062,401 bp or 2.0% of Mm10. In other words, the 5’-to-3’ span of mouse genes on the RefGene list should capture all of the RNA-seq true positives and almost 60% (2.0/3.4 × 100%) of the cCREs. With most cCREs overlapping annotated mouse genes, we can directly compare FFPE-CUTAC fragment counts to RNA-seq fragment counts by asking how well they correlate with one another over genes. Whereas FFPE-CUTAC replicates and RNA-seq replicates are very strongly correlated to a similar extent, with “arrowhead” scatterplots (R^2^ = 0.955-0.997), comparisons between FFPE-CUTAC and RNA-seq samples are “fuzzy” but nevertheless show strong correlations (R^2^ = 0.764-0.881) (**Figure 7a**).

**Figure 7:**
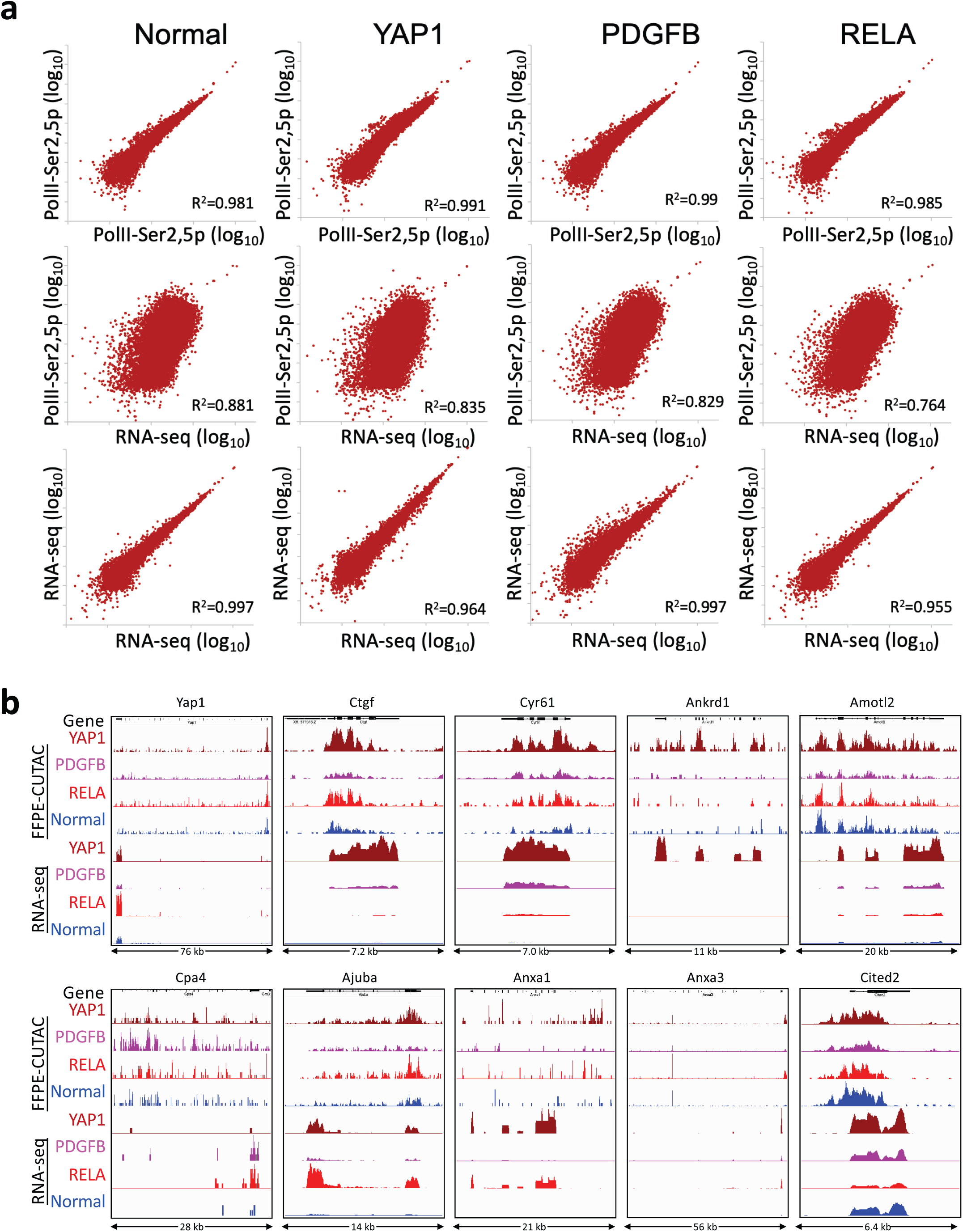
Comparisons between FFPE-CUTAC and RNA-seq. **a**) Top panels: Scatterplots of representative FFPE-CUTAC replicate samples from RNAPII-Ser2,5p for normal brain and the three tumorour brains. Middle panels: Scatterplots of comparisons between the RNAPII-Ser2,5p sample and the corresponding RNA-seq dataset. Lower panel: Scatterplots of RNA-seq datasets used in the comparisons. **b**) Comparisons between FFPE-CUTAC and RNA-seq over Yap1 and previously reported Yap1 direct targets. Tracks were group-autoscaled within modalities.

We also determined the extent to which the same genes differ significantly between tumor normal in the two datasets. Using an FDR = 0.05 cut-off for both FFPE-CUTAC and RNA-seq, we found that 80-82% of genes were found in both lists: 52 of 63 for Yap-driven tumors versus normal brains, 268 of 336 for PDGFB-driven versus normal and 1519 of 1896 for RELA-driven versus normal. However, there is a striking difference in the specificity with which these genes are identified as illustrated by comparison of volcano plot displays: FFPE-CUTAC provides high specificity for regulatory elements, where significant differences between cCREs are almost exclusively at the upregulated corner of the volcano plots (high positive log^2^ fold-change, high —log^10^ FDR) (**Figure 5**). In contrast, ∼1/3 to 1/2 of 23,551 genes show significant differences between these tumorous and normal brains using RNA-seq with massive, mostly symmetrical “volcanic eruptions” (**Figure S2**).

To validate these comparisons, we aligned profiles of FFPE-CUTAC and RNA-seq at YAP1 and at nine direct targets of YAP1, which were previously determined based in part on the RNA-seq data (43). As expected, the FFPE-CUTAC profiles are enriched primarily at 5’ ends and RNA-seq at 3’ ends (**Figure 7b**). Importantly, all ten examples showed full or partial concordance between FFPE-CUTAC and RNA-seq. We conclude that there is overall excellent agreement between our FFPE-CUTAC data and previously published high-quality RNA-seq datasets. The very high specificity of FFPE-CUTAC data, together with its simple implementation and potential for automation, make it a unique and potentially useful modality for discovery of functional biomarkers.

## Discussion

Fixation-related DNA and chromatin damage has thus far impeded the practical application of chromatin profiling to FFPEs (4). Here we have shown that improvements to the single-tube CUT&Tag-direct protocol to make it suitable for whole cells, and together with heat-treatment of deparaffinized needle-extracted 10-micron FFPE sections, provides high-quality CUTAC data. By using an RNAPII-Ser5p antibody for paused RNAPII, our FFPE-CUTAC data provides a ground-truth interpretation of accessibility, applicable to both promoters and enhancers (44). While RNA-seq has been the preferred method for profiling the transcriptome, it is strongly biased towards abundant transcripts, whereas transcription factors that drive development and are deregulated in cancer may be expressed at relatively low levels and can be difficult to detect. In contrast, the 343,731 genomic sites annotated as candidate *cis*-regulatory elements in the mouse genome can potentially provide direct information on transcriptional regulatory networks. Remarkably, nearly all significant differences between tumors and normal brain corresponded to increases in RNAPII and H3K27ac, a histone mark of active promoters and enhancers. Global hypertranscription is a general feature of aggressive human cancers (24), and the much better discrimination of RNAPII that we observed for FFPE-CUTAC over cCREs than for high-quality RNA-seq data over genes encourages more general application of FFPE-CUTAC technology for diagnosis, biomarker discovery and retrospective studies.

Cross-links and adducts resulting from the long incubations in formaldehyde necessary for long-term preservation cause DNA breaks and lesions that are serious impediments for most genomic methods applied to FFPEs. Indeed, standard CUT&Tag failed for the group that developed FACT-seq (16), and we also failed to obtain usable profiles for repressive H3K27me3 and H3K9me3 and gene-body H3K36me3 histone epitopes. We attribute these failures to the tight wrapping of DNA to lysine-rich histones, which are the most susceptible to cross-linking and formation of DNA adducts that result in DNA breaks during high-temperature cross-linking reversal (20). In contrast, nucleosome-depleted regions (NDRs) that are mapped using accessibility methods such as ATAC-seq and CUTAC are much better suited for FFPEs, as the protein machineries that occupy these sites are not especially lysine-rich. In particular, the YSPTSPS heptamer present in 52 tandem copies on the C-terminal domain of the largest subunit of RNAPII presents abundant lysine-free epitopes for CUT&Tag, and the use of low-salt tagmentation after stringent washes allows for tight binding of the Tn5 transposome within the confines of the NDR. We have previously shown that for epitopes such as H3K4 methylations (18) and RNAPII epitopes (45) that flank gaps in the nucleosome landscape at promoters and enhancers, tagmentation preferentially releases subnucleosomal fragments. FACT-seq improves yield with *in vitro* transcription from a T7 promoter inserted at single sites, however this strategy foregoes the advantage of the small size of NDRs at promoters and enhancers where nevertheless two Tn5s can fit with enough DNA in between for sequence-based mapping. We might attribute the better data quality that we obtained using CUTAC relative to FACT-seq to the very low probability of two Tn5s inserting close enough to one another and correctly oriented to produce a small amplifiable fragment by random chance. Curiously, H3K27ac FFPE-CUTAC detected cCREs even more sensitively than standard H3K27ac CUT&Tag on frozen tissue, which might indicate that better reversal of cross-links at NDRs than at nucleosomes facilitates tagmentation within NDRs while nucleosomes remain relatively intractable. Indeed, by avoiding the use of degradative enzymes and using only heat to expose epitopes in a suitable buffer, we found that bead-bound tissue shards from sheared FFPEs are much easier to handle without damage than cells or nuclei, where lysis and sticking is a constant concern.

We also discovered that DNA from *Rhodococcus erythropolis*, a species of bacteria that can live on paraffin wax as its only carbon source, is abundant in the FFPE samples that we processed, and this unfixed DNA competes against formalin-damaged DNA from FFPEs during PCR. As a result, lowering the amount of tissue in a paraffin slice results in relative increases in PCR products from *Rhodococcus* and other bacterial contaminants. To minimize the contribution of bacterial contaminants, we modified our single-tube CUT&Tag-direct method to use thermolabile Proteinase K prior to PCR, thus reducing cellular material that evidently inhibits PCR when using whole cells or FFPEs. As CUT&Tag-direct has been fully automated (25), we expect that our FFPE-CUTAC protocol will be suitable for institutional core facilities and commercial services, maximizing reproducibility and minimizing costs.

In conclusion, we have shown that RNAPII and H3K-27ac chromatin profiling can be conveniently and inexpensively performed on FFPEs in single PCR tubes. We use only heat in a suitable buffer to reverse the cross-links while making the tissue sufficiently permeable, followed by needle extraction and a slightly modified version of our CUT&Tag-direct protocol, which is routinely performed in many laboratories (19, 46). We found that data quality using low-salt tagmentation for antibody-tethered chromatin accessibility mapping is sufficient to distinguish cancer from normal tissues and resolve closely similar brain tumors. Using FFPE-CUTAC, our study identified direct targets of cancer drivers in tumors, validating our approach.

## Methods

### Cell lines

Human female K562 chronic myelogenous leukemia cells (American Type Culture Collection (ATCC)) were authenticated for STR, sterility, human pathogenic virus testing, mycoplasma contamination and viability at thaw. H1 (WA01) male hESCs (WiCell) were authenticated for karyotype, STR, sterility, mycoplasma contamination and viability at thaw. K562 cells were cultured in liquid suspension in IMDM (ATCC) with 10% FBS added (Seradigm). H1 cells were cultured in Matrigel (Corning)-coated plates at 37 °C and 5% CO^2^ using mTeSR-1 Basal Medium (STEMCELL Technologies) exchanged every 24 hours. K562 cells were harvested by centrifugation for 3 minutes at 1,000g and then resuspended in 1× PBS. H1 cells were harvested with ReleasR (STEMCELL Technologies) using the manufacturer’s protocols.

### Mouse tumor and normal tissues and FFPEs

Ntva;cdkn2a-/-mice were injected intracranially with DF1 cells infected with and producing RCAS vectors encoding either PDGFB (21), REL A-ZFTA (22), or YAP1-FAM118b (23) as has been described (47). When the mice became lethargic and showed poor grooming, they were euthanized and their brains removed and fixed at least 48 hours in Neutral Buffered Formalin. Tumorous and normal brains were sliced into five pieces and processed overnight in a tissue processor, mounted in a paraffin block and 10-micron sections were placed on slides. Slides were stored for varying times between 1 month to ∼2 years before being deparaffinized and processed for FFPE-CUTAC. Healthy mouse liver or intrahepatic cholangiocarcinomas tumors harvested from orthotopic models of intrahepatic cholangiocarcinoma mice with activating mutations of Kras^G12D^ and deletion of p53 (48) were fixed in formaldehyde for 7d before being sent to the Fred Hutch Experimental Histopathology Shared Resource for FFPE processing.

Deparaffinization was performed in Coplin jars using 2-3 changes of histology grade xylene over a 20-minute period, followed by 3-5 minute rinses in a 50:50 mixture of xylene:100% ethanol, 100% ethanol (twice), 95% ethanol, 70% ethanol and 50% ethanol, then rinsed in deionized water. Slides were stored in distilled deionized water containing 0.02% sodium azide for up to 2 weeks before use.

### Antibodies

Primary antibodies: H3K4me3: Active Motif cat. no. 39159; H3K27ac: Abcam cat. no. ab4729; RNAPII-Ser5p: Cell Signaling Technologies cat. no. 13523; RNAPII-Ser2,5p: Cell Signaling Technologies cat. no. 13546; H3K27me3: Cell Signaling Technologies cat. no. 9733; H3K4me2: Epicypher cal. no. 13-0027; H3K36me3: Thermo cat. no. MAS-24687. Secondary antibody: Guinea pig α-rabbit antibody (Antibodies on-line cat. no. ABIN101961).

### CUT&Tag-direct for whole cells

Concanavalin A (ConA) coated magnetic beads (Bangs Laboratories, ca. no. BP531) were activated just before use with Ca^++^ and Mn^++^ as described (19). Frozen whole-cell aliquots were thawed at room temperature, split into PCR tubes and 5 μL ConA beads were added with gentle vortexing. All subsequent steps through to library preparation and purification f ollowed t he standard CUT&Tag-direct protocol (19) using pAG-Tn5 (Epicypher cat. no. 15-1117), except that 1) All buffers from antibody incubation through tagmentation included 0.05% Triton-X100; 2) The fragment release step was performed in 5 μl 1% SDS supplemented with 1:10 Thermolabile Proteinase K (New England Biolabs cat. no. P8111S) at 37^°^C 1 hr followed by 58^°^C 1 hr; 3) SDS was quenched by addition of 15 μl 6% Triton-X100. A detailed step-by-step protocol is available at Protocols. io: https://www.protocols.io/view/cut-amp-tag-direct-for-whole-cells-with-cutac-x54v9mkmzg3e/v4, with a comment box for help.

### FFPE-CUTAC

Tissue sections on deparaffinized s lides w ere diced using a razor and scraped into a 1.7 mL low-bind tube containing 400 μl 800 mM Tris-HCl pH8.0, 0.05% Triton-X100. Incubations were performed at 80-90^°^C for 8-16 hours or as otherwise indicated either in a heating block or divided into 0.5 mL PCR tubes after needle extraction. Needle extraction was performed either before or after Concanavalin A-bead addition using a 1 ml syringe fitted with a 1” 20 gauge needle with 20 up-and-down cycles, and in some cases was followed by 10 cycles with a 3/8” 26 gauge needle. Other steps through to library preparation and purification followed the standard CUT&Tag-direct protocol (19) with the following exceptions: 1) All buffers from antibody incubation through tagmentation included 0.05% Triton-X100; 2) The fragment release step was performed in 5 μl 1% SDS supplemented with 1:10 Thermolabile Proteinase K (New England Biolabs cat. no. P8111S) at 37^°^C 1 hr followed by 58^°^C 1 hr; 3) SDS was quenched by addition of 15 μl 6% Triton-X100; 4) PCR was performed with an extension step (10 sec 98^°^C denaturation, 30 sec 63^°^C annealing and 1 min 72^°^C extension for 12 cycles). A detailed step-by-step protocol, including xylene and mineral oil deparaffinization options, is available on Protocols.io: <u>DOI: dx.doi.org/10.17504/protocols.io.14egn292zg5d/ v1</u>, with a comment box for help.

### DNA sequencing and data processing

Libraries were sequenced on an Illumina NextSeq 2000 instrument with paired end 50×50 reads. Adapters were clipped by cutadapt version 4.1 (49) with parameters -j 8 --nextseq-trim 20 -m 20 -a AGATCGGAAGAGCA-CACGTCTGAACTCCAGTCA -AAGATCGGAAGAGC-GTCGTGTAGGGAAAGAGTGT -Z Clipped reads were aligned by Bowtie2 version 2.4.4 (50) to the Mus musculus mm10 and Homo sapiens hg19 reference sequences from UCSC (51) and to the UCSC *Rhodococcus erythropolis* complete genome (NZ_CP007255.1) from NCBI with parameters --very-sensitive-local --soft-clipped-unmapped-tlen --dovetail --no-mixed --no-discordant -q --phred33 -I 10 -X 1000

### Data analysis

BLASTN searches of unmapped reads against the Nucleotide database were done on the NCBI web site (https://blast.ncbi.nlm.nih.gov/Blast.cgi?PRO-GRAM=blastn&PAGE_TYPE=BlastSearch&LINK_LOC=blasthome). We noticed the majority hit several bacteria so we narrowed the search to the RefSeq Genome Database restricted to Bacteria (taxid:2). After further analysis of these BLAST hits we made a Bowtie2 reference sequence from five bacteria:

NZ_CP007255.1 *Rhodococcus erythropolis* R138

NZ_JACNZU010000010.1 *Bacillus pumilus* strain 167T-6

NZ_JAGEKP010000001.1 *Leifsonia* sp. TF02-11

NZ_QCYC01000100.1 *Vibrio vulni icus* strain Vv003

NZ_MLHV01000015.1 *Mycobacterium syngnathi-darum* strain 24999

Properly paired reads were extracted from the alignments by samtools version 1.14 (45) bamtobed command into mapped fragment bed files and normalized count tracks were made by bedtools version 2.30 (52, 53) genomecov command with scale (size_of_reference_sequence/total_counts). Normalized count tracks are the fraction of counts at each base pair scaled by the size of the reference sequence so that if the counts were uniformly distributed across the genome there would be one at each position. Distributions of the lengths of the mapped fragments were made using the UNIX sort and uniq -c command. Peaks were made by MACS2 version 2.2.6 (54) from the mapped fragment bed files with parameters:

macs2 callpeak -t <fragments> -f BEDPE -g hs --keep-dup all -p 1e-5 -n <name> --SPMR For comparisons, the following datasets were downloaded from GEO: GSM5530653-55 (mouse kidney H3K27ac FACT-seq replicates 1-2 and H3K27ac Frozen CUT&Tag, respectively), GSM5530669-70 (mouse liver H3K27ac FACT-seq replicates 1-2) and GSE172688 (ENCODE ChIP-seq mouse post-natal forebrain).

Random sub-samples of fixed sizes were taken from the mapped fragment bed files using the UNIX shuff command and peaks were found by MACS2 for each sub-sample. Then the fraction of reads in peaks (FRiP) was computed using the bedtools intersect command. Although sequencing data from Reference 16 (GEO GSE171758) was single-end yielding median fragment lengths of 51 bp for kidney and 75 bp for liver, these fragment lengths were sufficiently similar to our paired-end median fragment lengths (65-76 bp for brain and 63-68 bp for liver) that no adjustments were made in comparisons.

cCRE overlaps were calculated for 10 million mapped fragments per sample as the number of fragments with at least one basepair overlap with a cCRE. Differential analyses of FFPE-CUTAC and RNA-seq data were performed using the Voom/Limma option (33) on the Degust server (https://degust.erc.monash.edu/).

Files for degust (https://degust.erc.monash.edu/) were made for a list of 343,731 Candidate *cis-*regulatory elements (cCREs) for *Mus musculus* from ENCODE (ENCFF427VRW) and for 23,551 genes from the Mus musculus Mm10 refGene list from the University of California Santa Cruz Genome Resource. The refGene file contains multiple transcripts for each gene so we winnowed it by using the region from the minimum start position to the maximum end position for each set of transcripts for a gene. For sums, we added the normalized counts within each cCRE or gene region for analysis by the degust web site. For summits we took the maximum within each region.

## Acknowledgements

We thank Christine Codomo, Doris Xu and Terri Bryson for technical assistance, Iris Luk for generating the cholangiocarcinoma model, Matthew Fitzgibbon for bioinformatics support, the Fred Hutch Genomics Shared Resource for sequencing and data processing and the Fred Hutch Experimental Histopathology Shared Resource for FFPE processing of liver samples.

## Author Contributions

S.H. and E.C.H. conceived the study; S.H. and R.M.P. performed the experiments; F.S., Z.R.R., S.K. and E.C.H. provided critical materials; D.H.J. advised on the methods; S.H. and J.G.H. analyzed the data; S.H. wrote the manuscript; S.H., J.G.H., K.A., S.K. and E.C.H. reviewed and edited the manuscript, and all authors approved the manuscript.

---

**Figure S1:**
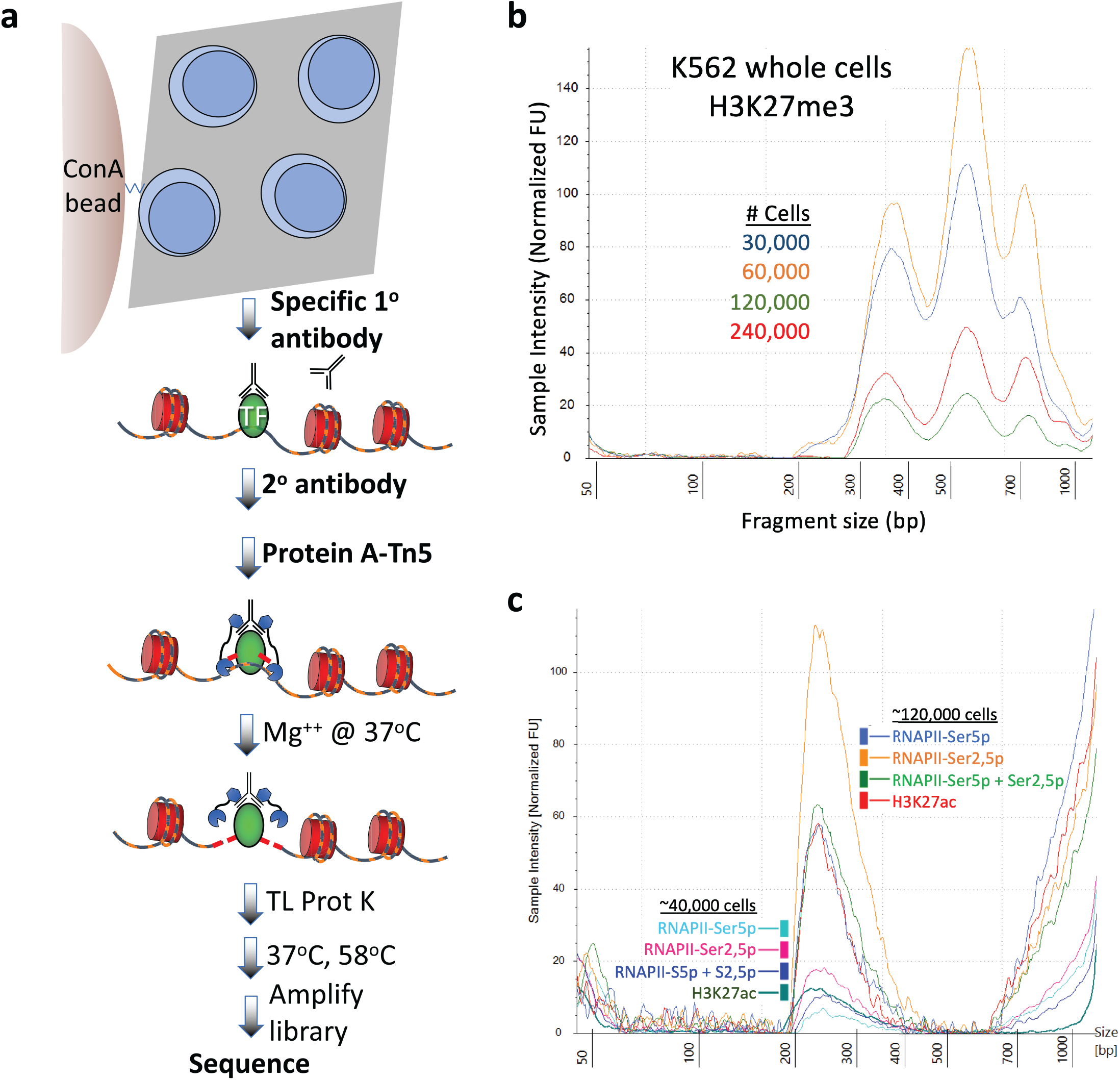
Modified CUT&Tag-direct for whole cells and FFPEs. **a)** Scheme, where TL Prot K is Thermolabile Proteinase K (New England Biolabs). **b**) Representative Tapestation profiles for whole-cell CUT&Tag-direct. A log culture of K562 cells was supplemented with 10% DMSO, concentrated to 2 million cells/ml, aliquoted, slow-frozen in Mr. Frosty containers and stored at -80 °C. An aliquot was thawed and 15-60 μL was dispensed into PCR tubes for CUT&Tag-direct using an H3K27me3 antibody (CST cat. no. 9733). **c)** Tapestation profiles for FFPE CUTAC samples pre-incubated at 85 ^°^C for 12 hr using four different antibodies. Each sample was divided 3/4-1/4 in the TAPS-wash before fragment release. Antibodies diluted 1:25 were RNAPII-Ser5p Cell Signaling Technology #13523, RNAPII-Ser2,5 Cell Signaling Technology #13546 and H3K27ac: Abcam #4729. A 10-micron section of a mouse brain tumor FFPE was deparaffinized using xylene. Note that both the CUTAC peaks the high-molecular weight smears scale with the amount of sample, likely representing ambient RNAs, which do not interfere with flow cell runs.

**Figure S2:**
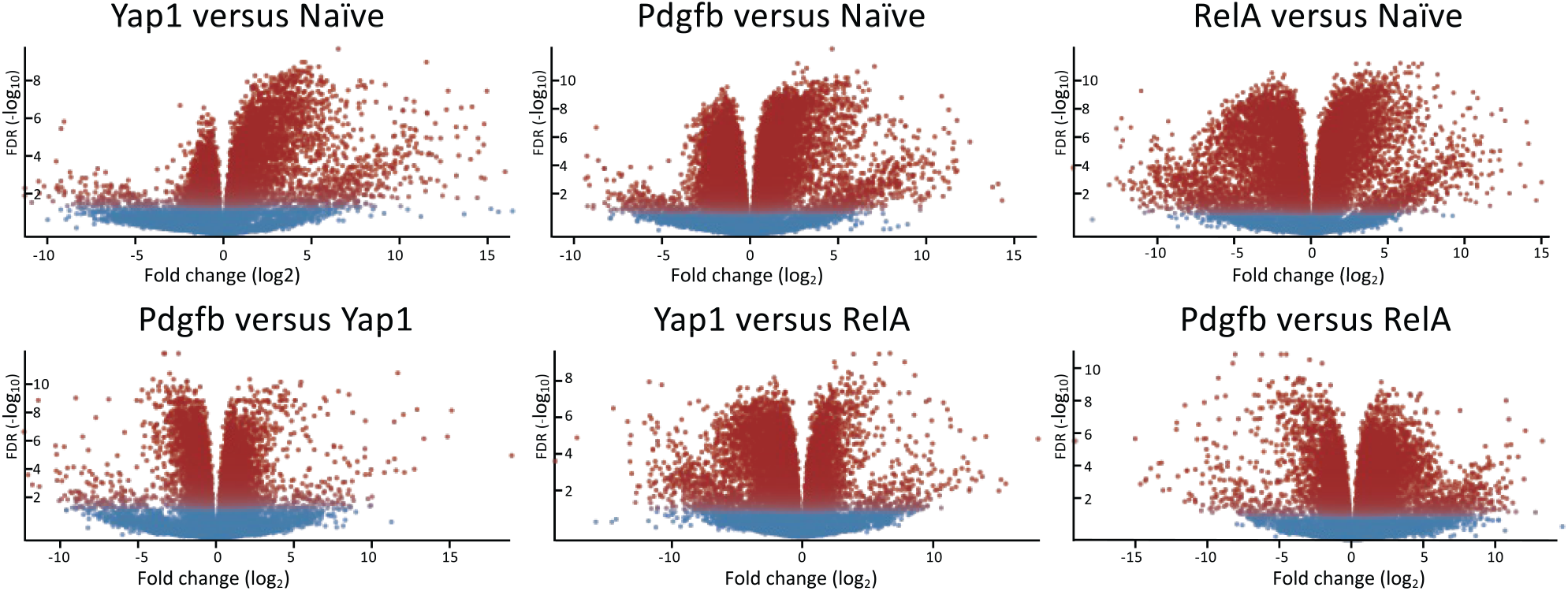
Volcano plots of RNA-seq comparisons. YAP1: 3 replicates; PDGFB: 4 replicates; RELA: 4 replicates; Normal: 7 replicates.

**Table S1:**
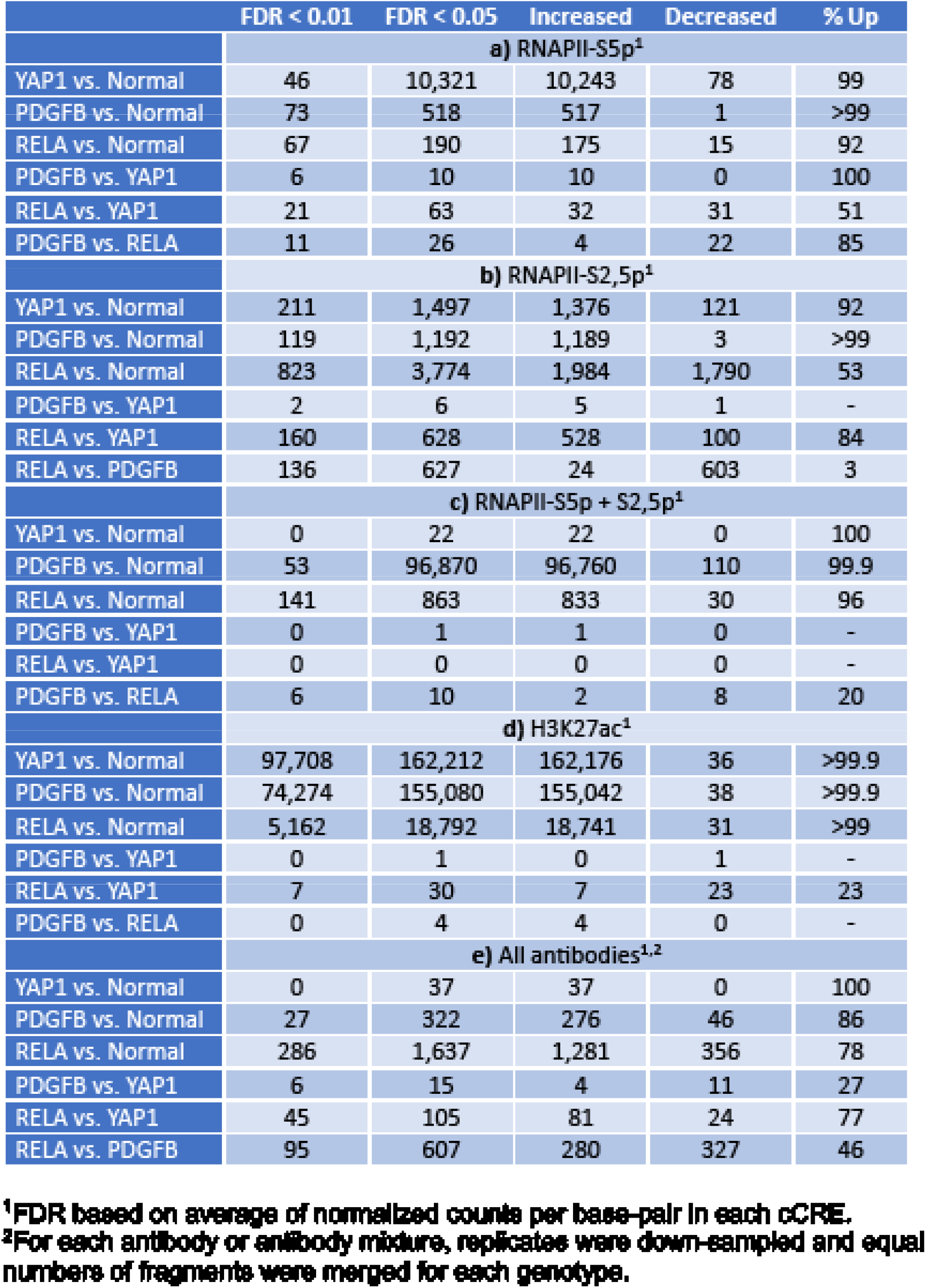
Signficant cCRE differences between FFTPE-CUTAC datesets

